# In cell architecture of the nuclear pore complex and snapshots of its turnover

**DOI:** 10.1101/2020.02.04.933820

**Authors:** Matteo Allegretti, Christian E. Zimmerli, Vasileios Rantos, Florian Wilfling, Paolo Ronchi, Herman K.H. Fung, Chia-Wei Lee, Wim Hagen, Beata Turonova, Kai Karius, Xiaojie Zhang, Christoph Müller, Yannick Schwab, Julia Mahamid, Boris Pfander, Jan Kosinski, Martin Beck

## Abstract

Nuclear pore complexes (NPCs) mediate exchange across the nuclear envelope. They consist of hundreds of proteins called nucleoporins (Nups) that assemble in multiple copies to fuse the inner and outer nuclear membranes. Elucidating the molecular function and architecture of NPCs imposes a formidable challenge and requires the convergence of *in vitro* and *in situ* approaches. How exactly NPC architecture accommodates processes such as mRNA export or NPC assembly and turnover inside of cells remains poorly understood. Here we combine integrated *in situ* structural biology, correlative light and electron microscopy with yeast genetics to structurally analyze NPCs within the native context of *Saccharomyces cerevisiae* cells under conditions of starvation and exponential growth. We find an unanticipated *in situ* layout of nucleoporins with respect to overall dimensions and conformation of the NPC scaffold that could not have been predicted from previous *in vitro* analysis. Particularly striking is the configuration of the Nup159 complex, which appears critical to spatially accommodate not only mRNA export but also NPC turnover by selective autophagy. We capture structural snapshots of NPC turnover, revealing that it occurs through nuclear envelope herniae and NPC-containing nuclear vesicles. Our study provides the basis for understanding the various membrane remodeling events that happen at the interface of the nuclear envelope with the autophagy apparatus and emphasizes the need of investigating macromolecular complexes in their cellular context.

## Text

Nuclear pore complexes (NPCs) are giant macromolecular assemblies with a very intricate architecture. About 30 different genes termed nucleoporins (Nups) encode for components of NPCs. Scaffold Nups contain folded domains and form a cylindrical central channel. This channel is lined with FG-Nups, which harbor intrinsically disordered FG-rich repeats that interact with nuclear transport receptors. Nups assemble in multiple copies to form an eight-fold rotationally symmetric complex, totaling in ~550 protein building blocks in yeast and ~1000 in mammals^1^. The NPC consists of two outer rings, also called nuclear (NR) and cytoplasmic rings (CR), which are placed distally to the inner ring (IR) that resides at the fusion plane between the inner and outer nuclear membranes. Within the three-ringed architecture, Nups are organized into subcomplexes that form specific substructures. The Y-complex is the key scaffolding component of the outer rings, whereas the IR scaffold is built by the inner ring complex^2^. Further, the yeast Nup159 complex (Nup214 complex in mammals) associates asymmetrically with the Y-complex at the CR and facilitates the terminal steps of mRNA export. Its core consists of two Nup159-Nup82-Nsp1 heterotrimers that dimerize into a characteristic P-shaped structure^3,4^.

Several studies have highlighted that the accurate spatial positioning of the Nup159 complex with respect to the central channel at the cytoplasmic face of the NPC is critical for the spatial organization and directionality of mRNA export^3,5,6^. While mRNAs are chaperoned through the FG repeats within the central channel in a Mex67-dependent (but Ran independent) manner, exported mRNPs encounter the ATPase activity of the DEAD-box RNA helicase Dpb5 at the cytoplasmic face that removes Mex67 and ratchets the RNA into the cytoplasm^1,6,7^. Dbp5 recruitment and positioning is ensured by the N-terminal beta propeller of Nup159, which is separated by a flexible linker from the C-terminal coiled coils that anchor Nup159 to the NPC scaffold^7^.

However, understanding how Nup subcomplexes are positioned to each other within the context of the nuclear membranes and the overall architecture of NPCs imposes a considerable challenge to structural biologists and requires the convergence of *in vitro* and *in situ* approaches^8^. While several high-resolution structures of Nups have been solved by X-ray crystallography, biochemical analysis and cross-linking mass spectrometry revealed the interaction topology. Cryo-electron microscopy (cryo-EM) maps provided an overall framework for the positioning of subcomplexes using systematic fitting approaches^9,10,11,12,13^. Cryo-electron tomography (cryo-ET) with subsequent subtomogram averaging is the method of choice to generate such cryo-EM maps, because it allows structural analysis of NPCs within the native context of the nuclear membranes that have to be considered as an integral part of their architecture. Such *in situ* structural analysis is available for both mammals and algae (reviewed in^1^) and revealed that key features of the NPC architecture, such as the stoichiometry of Y-complexes within the outer rings, are not conserved. Intriguingly, the P-shaped outline of the yeast Nup159 complex^4^ was not apparent in any of the cryo-EM maps available to date, either suggesting structural diversity, or alternatively, questioning its physiological relevance.

To date, *in situ* structural analysis is still missing for the *S. cerevisiae* NPC (*Sc*NPC), which has been extensively studied as a model organism not only for Nup structure, NPC architecture and the mechanism of mRNA export, but also NPC surveillance and turnover^1,14^. Thus far, only detergent extracted, purified NPC species have been analyzed^13^ that bear neither the nuclear membranes nor any distinct features of Y-complexes (reviewed in^1^). An integrative model of the entire *Sc*NPC architecture based on extensive experimental analysis *in vitro* has been recently put forward^13^ and in principle could enable structure-to-function analyses based on the very powerful yeast genetics approaches. However, it remains unknown to what extent the native architecture of the *Sc*NPC has been preserved *in vitro*. Therefore, there has been a pressing need to elucidate the spatial organization of subcomplexes of the *Sc*NPC *in situ* within the cellular context.

### Structural analysis of the ScNPC in the cellular context

To characterize *Sc*NPC architecture *in situ*, we prepared thin cryo-FIB lamellae of exponentially growing, plunge frozen cells^15^. We acquired 230 cryo-electron tomograms (Fig. 1a) containing ~500 *Sc*NPCs and determined the structures by subtomogram averaging (see Methods). The resulting cryo-EM map at ~25 Å resolution (Extended Data Fig. 1a-b) provides the most detailed overview of the native configuration and conformation of subcomplexes of actively transporting NPCs in cells of any species to date (Fig. 1b-c, Extended Data Fig. 1b-c). A visual inspection of the structure reveals striking disparities, not only compared to the previously analyzed structures of the algal NPC^10^ and human NPC^9^ that likely account for species-specific differences, but also in comparison to previous cryo-EM maps of the biochemically isolated and detergent-extracted *Sc*NPC (Extended Data Fig. 1c-d, Source Data 1 attached for review process). The previously published integrative model of one asymmetric unit of the IR^13^ fits unambiguously into the observed density (p-value 1.6*10^-12^ Extended Data Fig. 2), however the model of the entire IR has to be dilated by ~20 nm in diameter in order to fit the *in situ* conformation (Extended Data Fig. 3a), thereby spatially separating the eight individual spokes (Extended Data Fig. 3a). This analysis underlines the plasticity of the NPC within cells^15^, which might be physiologically relevant for the transport of large cargos and inner nuclear membrane proteins.

**Fig. 1:**
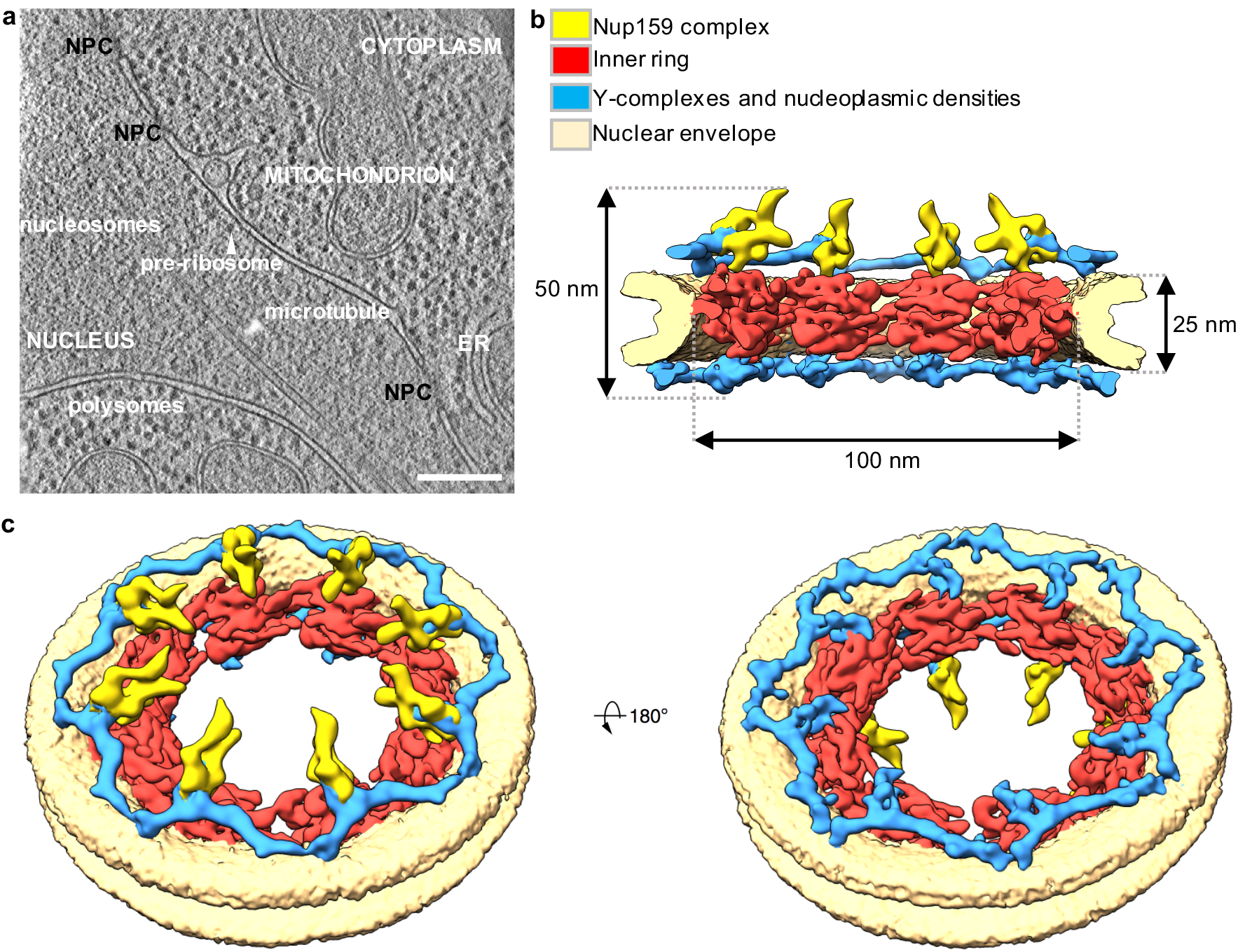
In cell structure of *S. cerevisiae* NPC. **a,** Tomographic slice of a *S. cerevisiae* cell during division. Scale bar 200 nm. **b**, Segmentation of the cryo-EM map of the *Sc*NPC cut in half, **c**, Tilted view of the entire NPC showing both the cytoplasmic (left) and the nucleoplasmic (right) face.

As expected^13,16^, 16 nuclear and cytoplasmic rings of Y-complexes are apparent in our cryo-EM map (Fig. 1c-d, Extended Data Fig. 1c), although some additional, yet unexplained density is observed in the nuclear ring. While the crystal structures and homology models of the yeast Y-complex vertex^17^ (Sec13–Nup145C–Nup84–*Sc*Nup120–*Sc*Seh1–*Sc*Nup85) or its fragments fit into the observed Y-shape density with significant p-values (Extended Data Fig. 2), the previous integrative model of the Y-complex^13^ does not fit due to the curvature of its tail (Extended Data Fig. 3c). We therefore applied an integrative modeling procedure (Methods) to take the observed *in situ* conformation into account and obtained complete structures of the Y-complexes in both CR and NR. Due to the larger circumference of the NPC, the Y-complexes occupy a more extended conformation (Extended Data Fig. 3b). These findings further emphasize that the combination of *in vitro* and *in situ* structural analysis is essential to understand NPC architecture in physiological conditions.

A prominent feature of the CR, which remains yet unassigned after localization of the Y-complexes, is the P-shaped density of the Nup159 complex that is strikingly reminiscent of the p
revious *in vitro* analysis of the isolated, negatively stained Nup159 complex^3,4^ (Fig. 2). Systematic fitting of the Nup159 complex negative stain map^4^ into our *in situ* structure confirmed this fit (Fig. 2b, p-value 0.0027, Extended Data Fig. 4a, see Methods). Based on the top resulting fit, we superimposed a previously published representative integrative model of the Nup159 complex^3,13^ and we locally fit it into our cryo-EM map (Extended Data Fig. 4b). In comparison to previous architectural models of the entire *Sc*NPC^3,13^, the P-shape is flipped around the axis that points towards the central channel (Extended Data Fig. 4b). Thereby, the Nup82 ß-propellers are positioned towards the inner ring (Extended Data Fig. 4b). The previous architectural model was in part guided by distance restraints from crosslinking mass spectrometry. In our updated Nup159 complex configuration, all four of the previously published crosslinks^13^ between the Nup159 and Y-complexes that could be mapped to the structure are satisfied (Extended Data Fig. 4e). The arm of the P-shape that consists of Nup159 DID tandem repeats binding multiple Dyn2 dimers^4,18^ is clearly apparent *in situ* (Fig. 2). It projects towards the cytoplasm at an angle of ~45° with respect to the nucleocytoplasmic axis (Fig. 2), while it had been previously thought to rather face the IR^3^ (Extended Data Fig. 4b). Instead, the protein termini preceding FG-repeats of Nsp1 face the central channel.

**Fig. 2:**
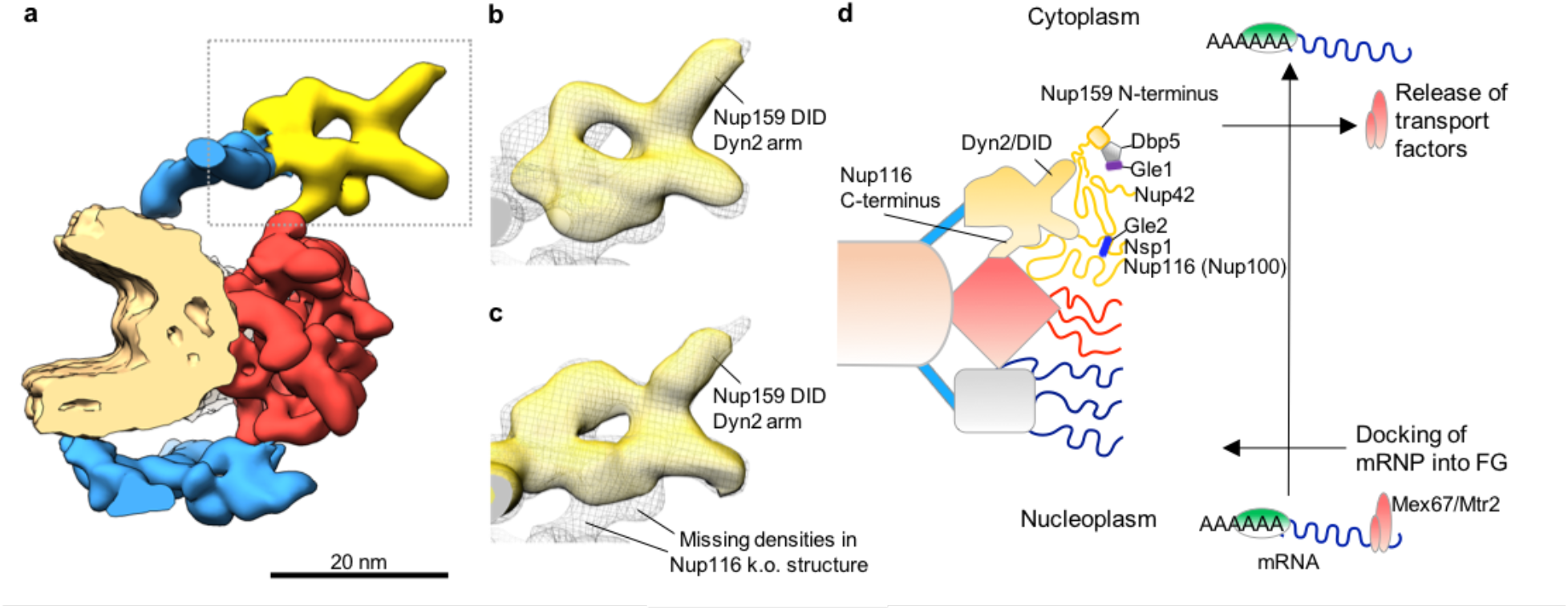
Nup159 complex architecture. **a**, Individual spoke of the *Sc*NPC (color-code as in Fig. 1). Nup159 complex is highlighted with a dotted frame. **b**, Nup159 complex region of the in cell *Sc*NPC map (gray mesh) superimposed with the top scoring systematic fit (see Methods) of the negative stain map of the Nup159 complex^4^ (yellow surface). **c**, Same as **b** but superimposed with the in cell *ScNPC* map of a *nup116Δ* strain grown at permissive temperature (yellow surface). **d**, Spatial model of how mRNA export is accommodated by the orientation of the Nup159 complex. A poly-A mRNA with transport factors Mex67/Mtr2 (red) and proteins that protect the poly-A tail from degradation (light green) docks to the unstructured region of the basket Nups^5^ (grey box with blue filaments). Nup159 N-terminal domain that mediates the release of the transport factors from the mRNA^7^ is anchored at the cytoplasmic side.

### Yeast genetics confirms orientation of the RNA export platform

To independently confirm this arrangement, we combined yeast genetics with *in situ* structural analysis. An important interactor of Nup82 is Nup116, one out of three yeast homologues of the essential vertebrate Nup98. This is a key Nup for establishing the NPC permeability barrier^19,20^, mRNA export^6,19^ and pre-ribosome translocation^21^. It is also an architecturally critical linker Nup that connects the inner ring, Y and Nup159 complexes through short linear motifs in its intrinsically disordered domains^12,22,23^. We superimposed the crystal structure of Nup116(966-1111)-Nup82(1-452)-Nup159(1425-1458)^24^ with the respective parts of two copies of Nup159 and Nup82 contained in the P-complex (Extended Data Fig. 4c-d). This analysis predicts that the autoproteolytic domain of Nup116 is placed into two yet unassigned densities that are proximate to each other and project towards Nup188 of the IR (Fig. 2c, Extended Data Figs. 4c-d and 5a). Crosslinking analysis^13^ and previous biochemical data suggesting that Nup116 links to Nup188 and Nup192 of the IR^22,23^ agree well with this configuration (Extended Data Figs. 4e and 5a). To validate this assignment, we structurally analyzed the *Sc*NPC in *nup116Δ* cells (knock-out, k. o.) cells at permissive temperature, at which NPCs have an ordinary morphological appearance in electron micrographs^25^. We acquired ~120 tomograms and obtained a cryo-EM map as described above (see Methods). The *nup116Δ* structure lacks density exactly at positions proximate to Nup188 as predicted by superposition of the Nup116-Nup82-Nup159 X-ray structure (Fig. 2c and Extended Data Figs. 4c and 5c). This finding corroborates our spatial positioning of Nup116 at the interface between the IR and Nup159-complexes and previous biochemical work that has established Nup116 as a linker nucleoporin associated with the inner ring components^22,23^. The positioning of the two copies of Nup116 agrees with only one of the two that were previously assigned by integrative modeling^13^ (Extended Data Fig. 5b). Our new model of NPC architecture spatiotemporally accommodates extensive biochemical analysis of mRNP export^5,7,19^. In fact, it places the FG-repeats that interact with Mex67 and are relevant earlier during the process, more towards the equatorial plane. Dbp5 and the N-terminus of Nup159 that facilitate the terminal release of Mex67 are positioned more towards the cytoplasm^26^ (Fig. 2d).

### The Nup159 complex mediates selective autophagy through nuclear envelope herniae

At nonpermissive temperature, the *nup116Δ* strain forms nuclear envelope (NE) herniae (Extended Data Fig. 6a), a phenotype in which NPCs are engulfed by the NE membranes and not accessible to the cytosol^25^. Interestingly, this phenotype was also observed when other Nup159 complex associated Nups, such as Gle2 were genetically perturbed^27^, and has been linked to the surveillance of NPC assembly by ESCRT proteins^28,29^. Since herniae were also observed under seemingly unrelated conditions, such as genetic perturbation of inner nuclear membrane proteins^30,31^ in both yeast and human cells^32^, their exact functional relevance remained to be determined^14,33^. We wondered if NPC architecture under the herniae is altered and addressed this using cryo-FIB milling, cryo-ET (~40 tomograms) and subtomogram averaging of NPCs at the herniae basis in the *nup116Δ* strain at non-permissive temperature. We found that in these trapped NPCs not only Nup116, but the entire cytoplasmic ring, including the Nup159 and Y-complexes are missing (Extended Data Fig. 6c). Concurrently, the nuclear envelope is less curved at the fusion plane, which is typical for the neck of herniae (Extended Data Fig. 6a-c). This result is in line with models that conceptualize herniae as failed inside-out assembly intermediates of NPCs, in which the lack of fusion of the NE membranes prevented the assembly of the cytoplasmic components (reviewed in^14,33^). Another finding further corroborates this model. We noticed that as compared to WT cells, the NE of *nup116Δ*. cells at permissive temperature contains a high number of mushroom-shape evagination of the inner nuclear membrane (Extended Data Fig. 6d-e). The structures are morphologically reminiscent of interphase assembly intermediates previously characterized in human cells^34^. These data point to a direct or indirect contribution of Nup116 to membrane fusion during NPC assembly.

A recent study identified Nup159 as an intrinsic receptor for selective autophagy of nuclear pores (Lee et al in revision; attached for review process). The AIM (Atg8-interacting motif), located proximate to the DID-Dyn2 arm of Nup159, mediates NPC turnover under conditions of low nitrogen supply. It recruits the ubiquitin-like autophagosomal protein Atg8 that is essential for anchoring autophagic protein cargo to autophagosomal membranes. Above findings suggest that the AIM is exposed into the cytoplasm together with the DID-Dyn2 arm (Fig. 3a), thus favoring the recruitment of cytoplasmic Atg8. We hypothesized that the herniae might be cleared by selective autophagy. This would make sense if they would not only occur upon genetic perturbation as previously observed, but also upon stress conditions that trigger autophagy, which was indeed the case. Under conditions of nitrogen starvation, NE herniae were observed, although they were less abundant as compared to the *nup116Δ* strain at nonpermissive temperature (Fig. 3b). How exactly herniae clearance could be mediated by the AIM of Nup159 in the *nup116Δ* background is however not obvious because Nup159 complex should not be present at the herniae according to the NPC structure determined in *nup116Δ* cells at not permissive temperature (Extended Data Fig. 6c).

**Fig. 3:**
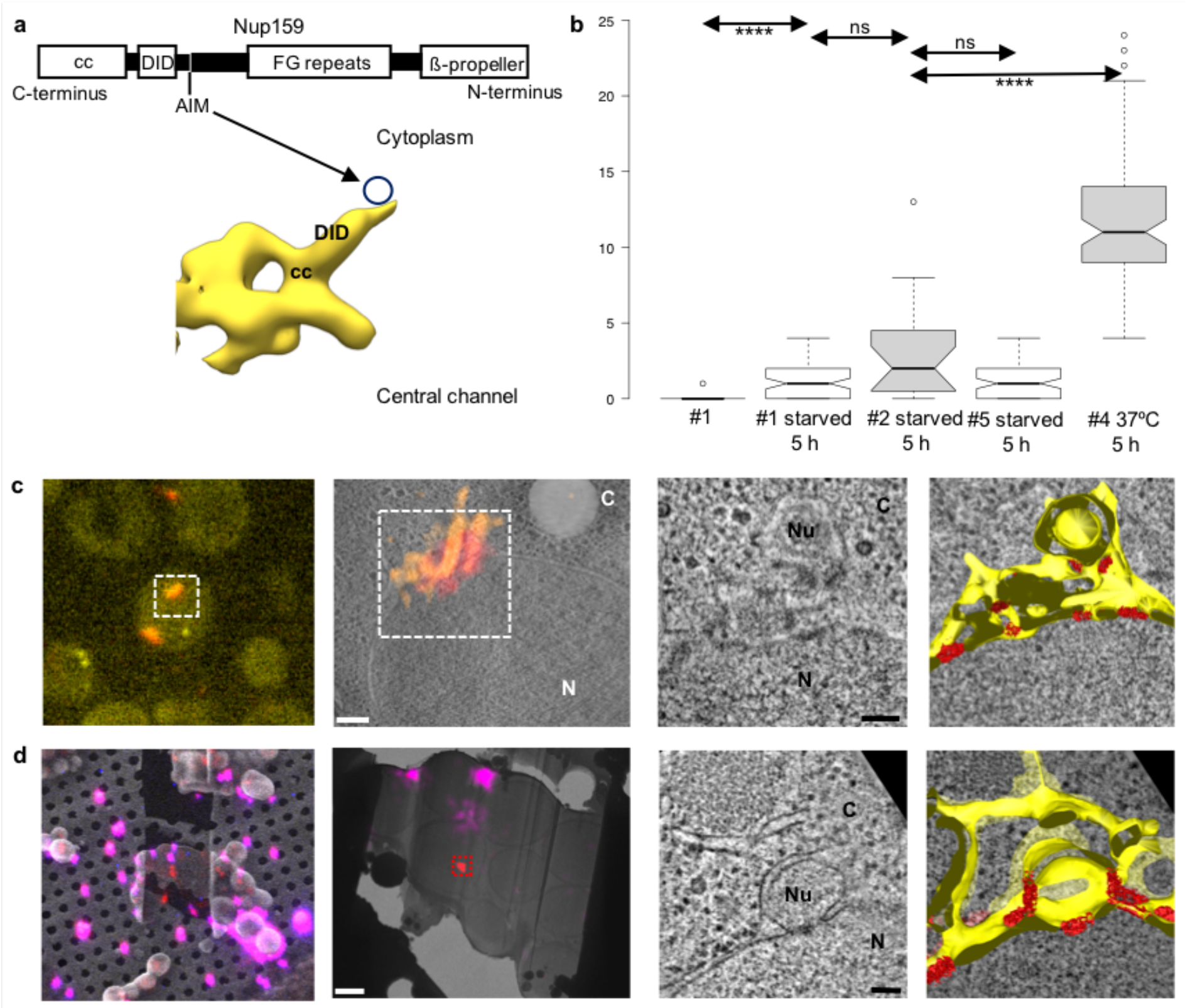
Cellular visualization of Nup159-Atg8 interaction by CLEM. **a**, Domain structure of Nup159 with corresponding map showing the position of the Atg8 interacting motif (AIM) (Lee et al). **b**, Box plot (median, 1st quartile) showing the number of herniae (see Methods) in *S. cerevisiae* strains before and after starvation according to Extended Data Table 1 (#1 control cells; #2 Nup159-Atg8-splitVenus, Nup170-Mars, *nup120D* cells; #4 *nup116Δ* cells; #5 *atg8Δ* cells). *nup116Δ* cells (#4) were shifted to the non-permissive Temperature of 37°C; herniae are induced (**** p<0.0001; Mann Whitney, two-sided). **c**, On section CLEM of strain #2 after 5.5h of starvation. The left panel shows the light microscopy signal on plastic section. SplitVenus signal arising from the interaction of Nup159 with Atg8 is shown in yellow. Nup170-Mars is shown in red. The second panel zooms in the area of interest (white dashed square in the previous panel) showing the overlay of the fluorescent signal on the tomographic EM slice. Scale bar: 200 nm. The framed area is shown as tomographic slice and segmented on the right (N nucleus; C cytoplasm; Nu nucleoplasm; scale bar: 100 nm; NPCs in red; membranes in yellow). **d,** 3D cryoCLEM of strain #2 after 5.5 h of starvation. SEM top view of a lamella with overlaid fluorescent signal. Red is NPC signal; magenta is fiducial-bead signal. Second panel: cryo-TEM overview (~2 nm/pix) of the same lamella. Scale bar: 2 μm. The framed area is shown as tomographic slice and segmented on the right. Scale bar: 100 nm.

To visualize how the autophagy machinery is recruited to the NPC, we employed a split-Venus approach specifically targeting the Nup159-Atg8 interaction. To enhance autophagic clearance of spatially clustered NPCs, we used a *nup120D* background^35^ as described in (Lee et al) but at shorter nitrogen starvation exposure (Fig. 3c-d and Extended Data Fig. 7a-b). We identified the regions of interest using correlative light and electron microscopy (CLEM) on plastic sections^36^ and 3D cryo-CLEM^37^. We acquired tomograms at positions of NPCs engaged in Atg8 interaction. These data revealed that agglomerates of herniae were engulfed by additional double membranes (Fig. 3c-d and Supplementary Video 1). Concurrently, the membrane topology of the herniae was more complex and displayed NPCs budding out of the NE, indicating additional membrane remodeling (Fig. 3c-d, Extended Data Fig. 7a and Supplementary Video 1). Thereby, many NPCs and thus AIMs are exposed to the cytoplasm (Fig. 3c-d, Supplementary Video 1). This finding would predict that herniae initially form independently from Atg8, possibly in the context of NPC surveillance^28^. Indeed, herniae formed in *atg8Δ* cells exposed to nitrogen starvation (Fig. 3b and Extended Data Fig. 8). We further noted, that under conditions where hernia formation was triggered but not selective autophagy, as in the *nup116Δ* strain at non-permissive temperature without starvation, herniae accumulate (Fig. 3b, Extended Fig. 6a) and are also observed as NPC-containing nuclear vesicles in the cytoplasm (Fig. 4c, Extended Data Fig. 9c and Supplementary Video 2), which agrees with the biochemical analysis of Lee et al Instead, under conditions in which selective autophagy is triggered by starvation, NPC-containing nuclear vesicles (~340 nm diameter) are surrounded by ribosomes and double membranes typical of autophagosomes (Fig. 4e-f and Extended Data Figs 7c and 9e-f), that are *en route* towards the vacuole as demonstrated by Lee et al (Extended Data Fig. 9g). Taken together, we observed several snapshots of NPC turnover, although the transitions between them, such as the budding of herniae into nuclear vesicles or fusion of autophagosome-engulfed NPC-containing-nuclear-vesicles to the vacuole need to be further investigated in the future (Extended Data Fig. 9). It will be also interesting to elucidate if herniae triggered under alternative genetic conditions, also in human cells^32^, ultimately feed into the autophagy pathway.

**Fig. 4:**
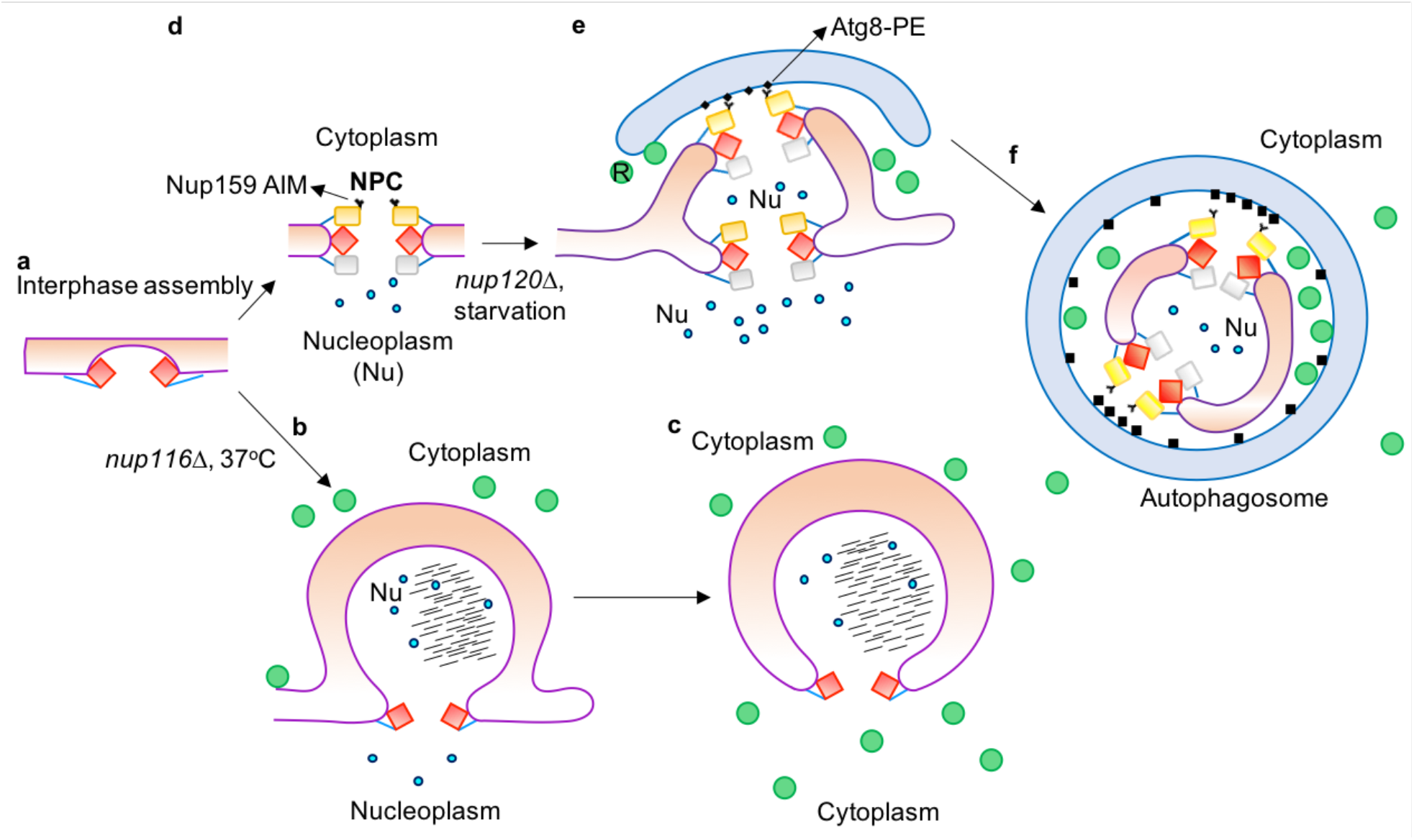
Cartoon model of membrane remodeling during NPC turnover by selective autophagy. **a,** Inside-out assembly intermediates^34^ or inner nuclear membrane evaginations (Extended Data Fig. 6d) progress into **b**, NE herniae as in the *nup116Δ* strain at non-permissive temperature (Extended Data Fig. 6a); or mature into **d,** fully assembled NPCs. **e**, When autophagy is triggered by Nitrogen starvation (or NPC clustering), NPCs are cytoplasmically exposed at the herniations by membrane remodeling (Fig. 3 c-d, Supplementary Video 1). Nup159 AIM interacts with Atg8 and double membranes are recruited to the nuclear envelope. **c**, If hernia accumulation is enforced in *nup116Δ* cells at non permissive temperature, nuclear vesicles are observed in the cytosol. **e, f**, If autophagy is triggered, nuclear vesicles are engulfed by double membranes (Extended Data Fig. 7c) and, as shown by Lee et al, ultimately end up in the lysosome (Extended Data Fig. 9g). Color-code as in Fig. 2d. R is ribosome depicted as a green circle; black lines represent dark densities visible in the cryo-tomograms.

In conclusion, our *in situ* structural analysis of *Sc*NPC clarifies the layout of the major Nup subcomplexes under native conditions. It confirms the physiological relevance of the P-shaped configuration of the Nup159 complex but identifies an unexpected orientation with respect to the scaffold that accommodates both the terminal steps of mRNA export (Fig 2d) and exposure of the intrinsic AIM (Fig. 4d). Our findings highlight the power of in cell structural biology to provide novel insights into fundamental processes of eukaryotic life.

---

**Extended Data Fig. 1:**
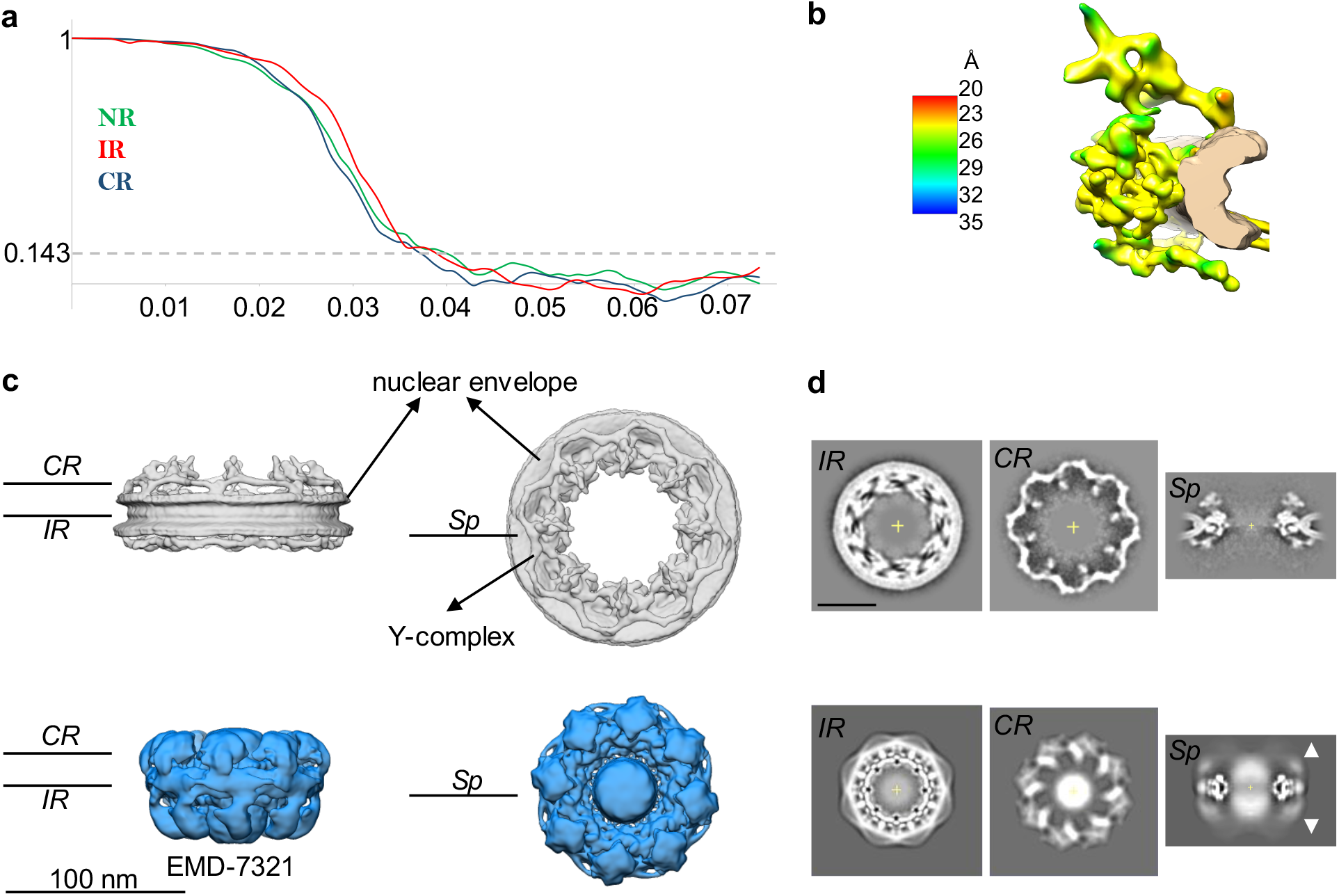
In cell structure of the *Sc*NPC vs detergent-extracted *Sc*NPC (EMD-7321). **a**, Gold standard FSCs of in cell cryo-EM map of the *Sc*NPC. All the curves (nuclear ring NR, cytoplasmic ring CR, inner ring IR) intersect the 0.143 criterium at around 25 Å resolution. **b**, Local resolution analysis^51^ with color-coded bar. **c**, In cell cryo-EM map (gray) in comparison to cryo-EM map of detergent-extracted *Sc*NPCs (blue, EMD-7321 at the suggested contour level) show significant differences in diameter and in interpretable features. The nuclear membranes and the Y-complexes are clearly discerned in the in cell cryo-EM map, in contrast to EMD-7321. **d**, Slices through the maps at the level of the cytoplasmic (CR) and inner rings (IR) and an individual spoke (Sp). Lines in c indicate slicing position shown in **d**. Arrowheads indicate blurred features in the outer rings. Scale bar: 50 nm. See also: Source Data 1 attached for review process.

**Extended Data Fig. 2:**
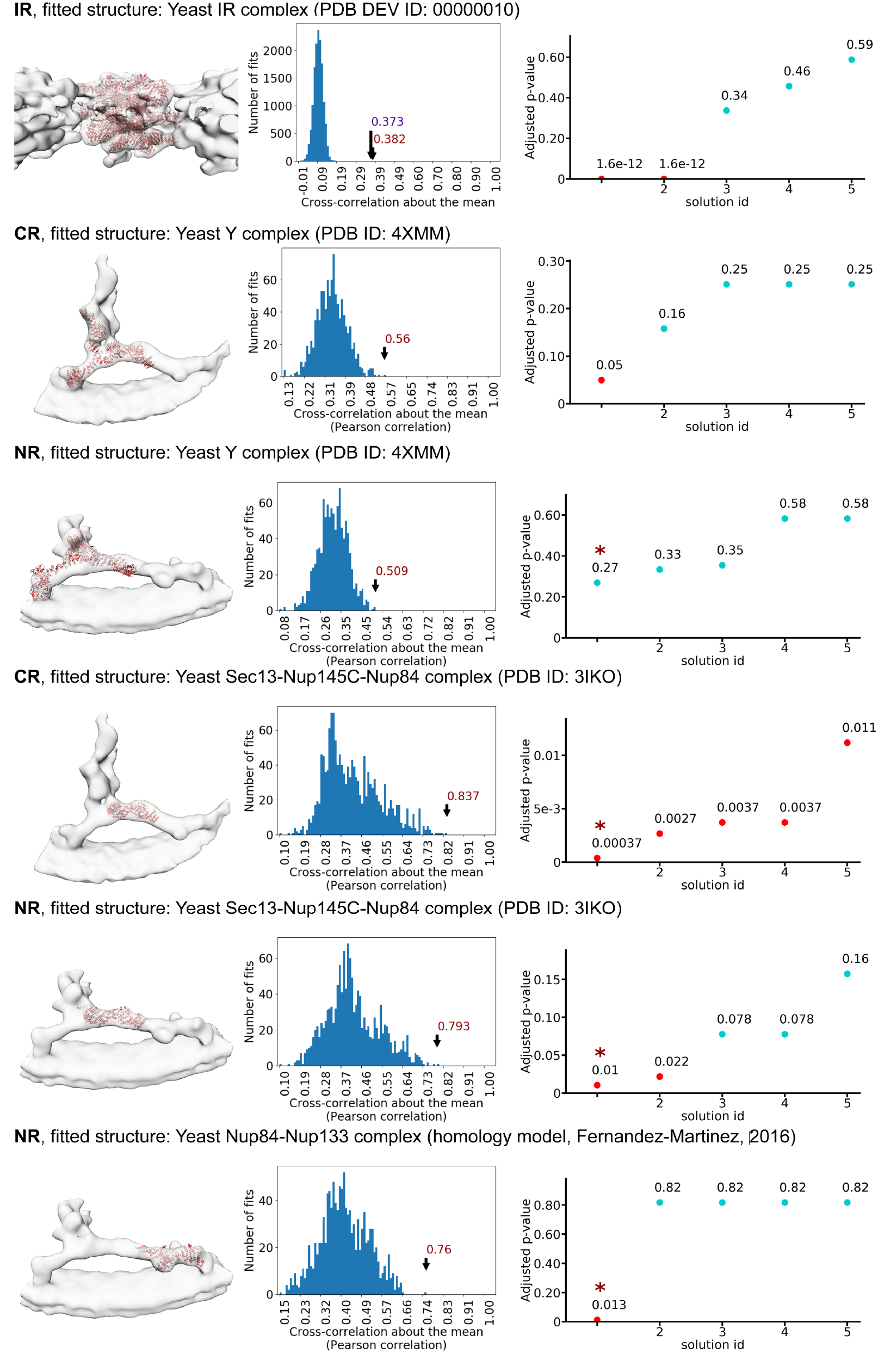
Systematic fitting of inner and outer ring components into the *Sc*NPC map. Each row shows the visualization of the most significant fits (left), the histogram of raw scores (middle), and a plot of the top five p-values (right). The statistically significant fits (p-value < 0.05, assessed as described in Methods) or the first top fit are marked with asterisk. The number of sampled fits used to calculate p-values after clustering of similar solutions was 14348, 1015, 1039, 1479, 1354, and 1183 for the rows from top to bottom. For the IR, the integrative model of the single spoke of the IR^13^ was used as the fitted structure. For the outer rings (CR and NR), the crystal structure of the yeast Y-complex was fitted or its parts corresponding to subcomplexes. The structures were fitted by an unbiased global search using UCSF Chimera^52^ and scored using the cross-correlation score about the mean as explained in the Methods. The IR complex was fitted to the entire spoke map, while the other structures were fitted to individual CR and NR segments.

**Extended Data Fig. 3:**
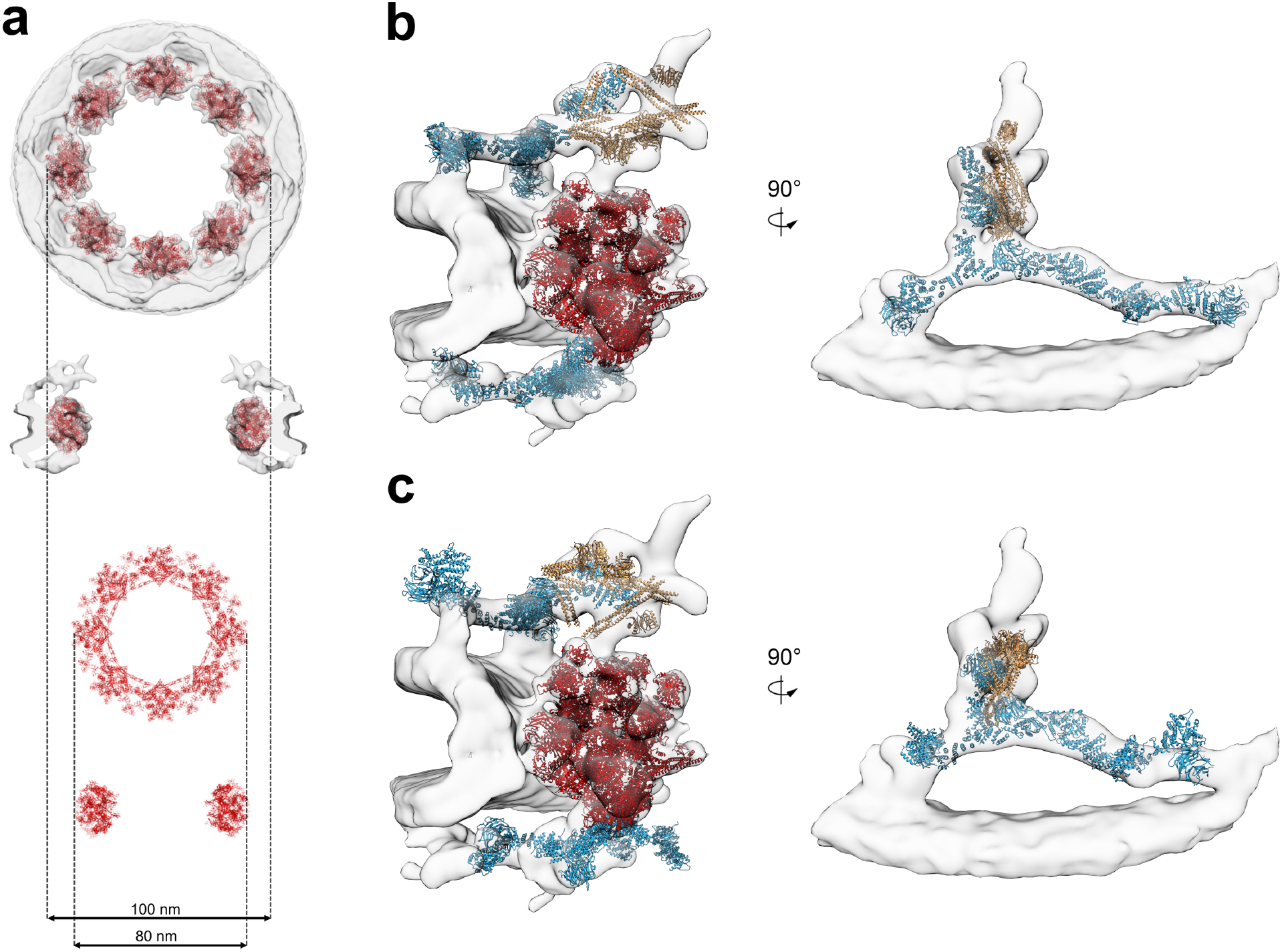
Architectural model of *Sc*NPC. **a**, Comparison of fitted (to the cryo-ET map from this study, depicted in gray) integrative Inner Ring complex (IRC) models (red ribbons) from^13^ with 20 nm diameter difference.**b**, Representative structural models of CR and NR Y-complex (blue ribbons), P-complex (yellow ribbons) and IR (red ribbons) built in this work (see Methods). The Y-complexes are more extended as compared to reference^13^ version by ~40 Å. **c**, Representative integrative structural models of CR and NR Y-complex (blue ribbons), P-complex (yellow ribbons) and inner ring complex (red ribbons) from^13^ fitted to cryo-ET map (grey density) from this study, with respect to spatial reference frame from^13^.

**Extended Data Fig. 4:**
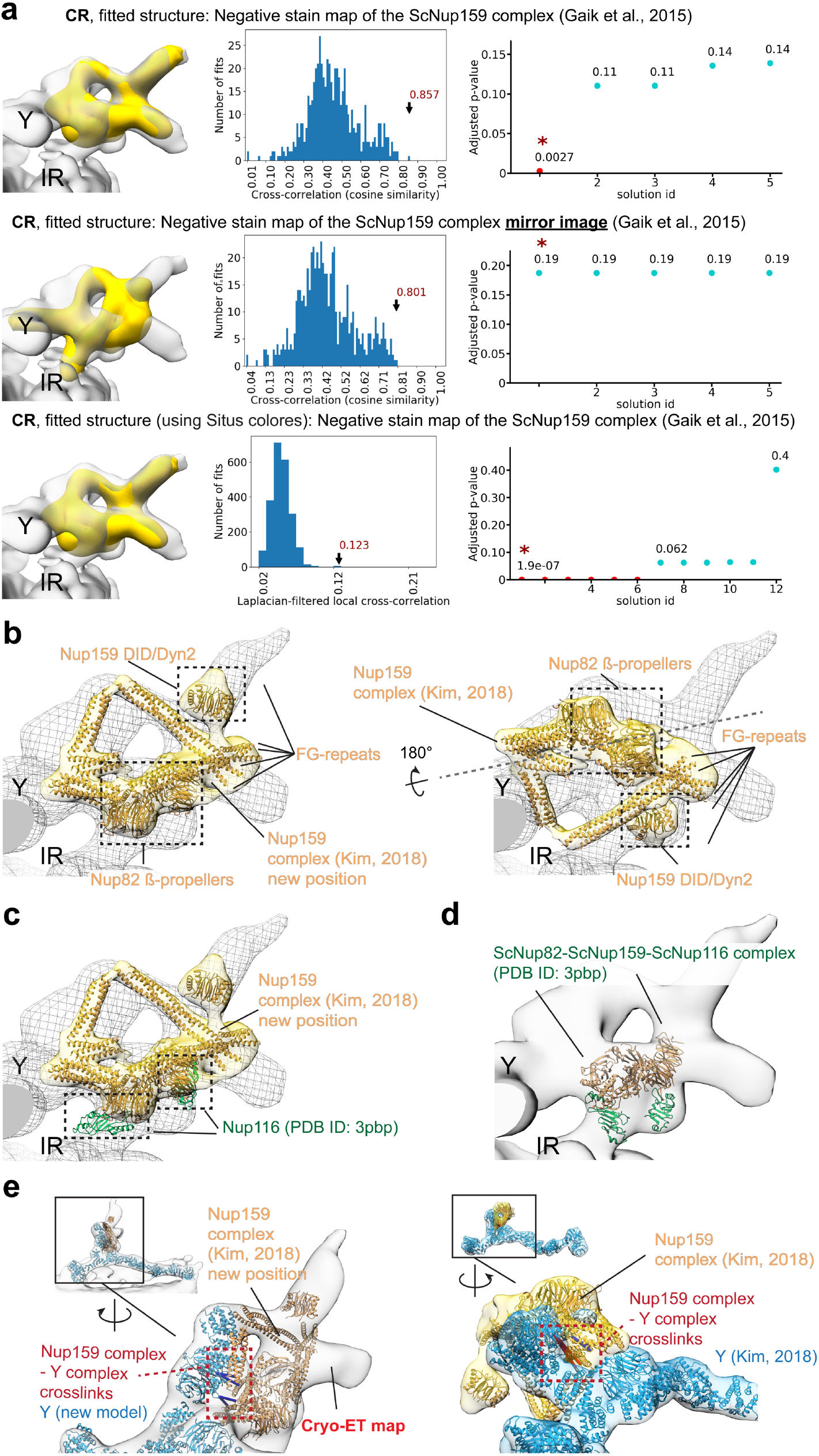
Validation of the orientation of the Nup159 complex. **a**, Systematic fitting of the negative stain map of the Nup159 complex into the *Sc*NPC map using UCSF Chimera^52^ and Colores program from the Situs package^54^. For comparison, both mirror images of the negative stain map (yellow) were fitted and only single one led to significant scores, underlining the unambiguous nature of the fit. Each row shows the visualization of the most significant fits (left), the histogram of raw scores (middle), and a plot of the top five p-values (right). The statistically significant fits (p-value < 0.05, assessed as described in Methods) or the first top fit are marked with asterisk. The number of sampled fits used to calculate p-values after clustering of similar solutions was 585, 599, and 2243, for top, middle, and bottom rows respectively. **b**, Representative integrative Nup159 complex model from^13^ inside the in cell *Sc*NPC map (gray mesh) in the orientation determined in this work (*left*) versus the previously published orientation (*right*). The Nup159 complex model is shown in orange ribbons within yellow localization probability density locally fitted (with UCSF Chimera^52^). The previous orientation was reproduced by first fitting the entire model from^13^ to the in cell cryo-EM map and then locally fitting the Nup159 complex to the density (which was needed to bring the Nup159 complex into the density and preserved the orientation). The dashed gray line indicates the flipping axis between the two fits. **c,** Superimposition of the crystal structure 3PBP^24^ onto representative integrative Nup159 complex model from reference^13^ in the newly determined orientation (left) predicts the position of Nup116, as confirmed by our knock-out study (Fig. 2c). **d,** Visualization of two of the top resulting systematic fits of the 3PBP crystal structure into the cryo-ET map presented in this study confirms our *nup116Δ* structure (Fig. 2c). **e**, Crosslinks between the Nup159 complex and the Y complex from^13^ support the new orientation (left) compared to the published orientation (right). Satisfied and violated crosslinks are depicted as blue and red bars, respectively.

**Extended Data Fig. 5:**
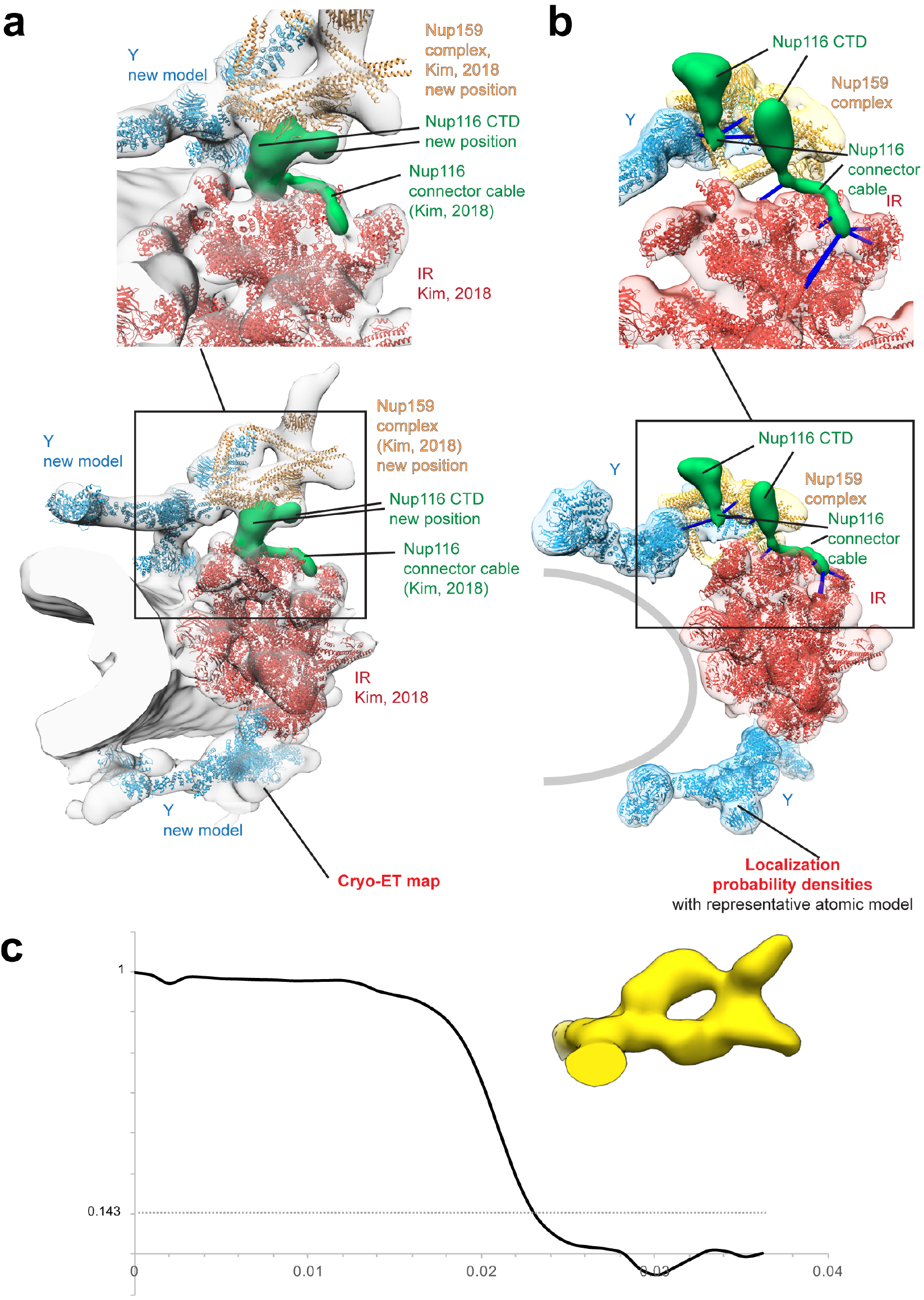
Nup116 positioning. **a**, New positioning of the Nup116 (left) versus previously published integrative model (PDB DEV ID: 00000010 from^13^, right). The two *Sc*NPC models were superimposed such that the IRs are aligned to the same reference frame. The Nup116 position is shown either as the density assigned to Nup116 based on the Nup116 knock-out structure (left) or as localization probability densities retrieved from reference^13^ model (right). The major structural elements of the NPC are indicated. The Nup116 connector cable in the in cell model (left) has been taken from reference^13^ based on its position relatively to the IR. Blue bars represent crosslinks from^13^ from Nup116 to other Nups. For the in cell model, the cryo-ET map is displayed; for the model from^13^, the localization probability densities (not an EM map) are shown instead. **b**, FSC of *nup116Δ* cytoplasmic filaments structure intersects the 0.143 line at ~40 Å, the cryo map is depicted in yellow as in Fig. 2c.

**Extended Data Fig. 6:**
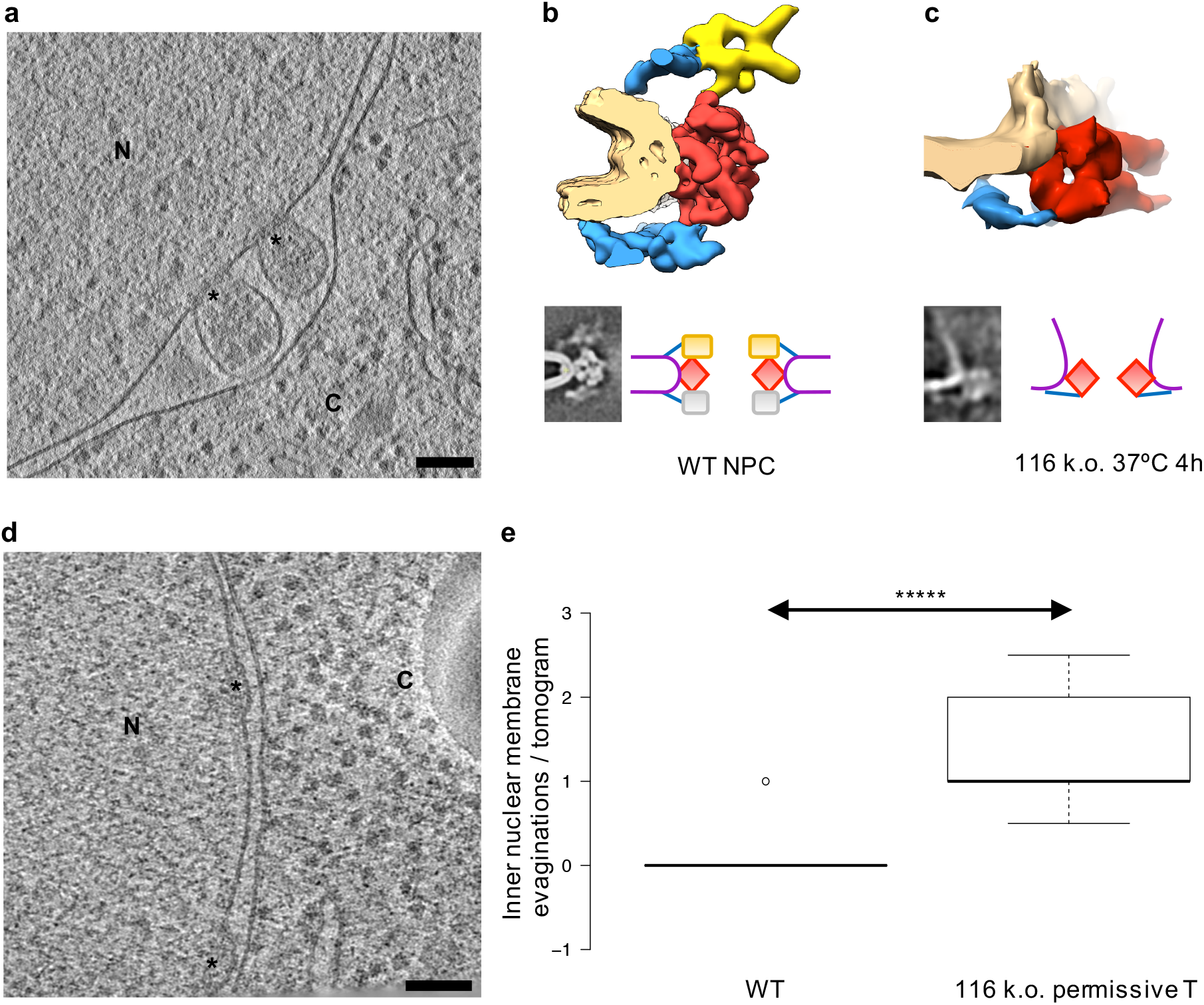
NPC and nuclear envelope morphology under *nup116Δ* conditions. **a,** tomographic slice of nuclear envelope with herniations (*nup116Δ* 4h at 37°C); N is nucleus, C is cytoplasm, stars depict density at the basis of the herniations that was subjected to subtomogram averaging; scale bar: 100 nm. **b,** Map and cartoon model of an individual spoke of the WT *Sc*NPC as in Fig. 2a and Fig. 4, shown isosurface rendered and as slice. **c,** same as **b** but as result of the subtomogram averaging at the basis of the herniations (see Methods). The average shows that the cytoplasmic ring, including the Nup159 complex is missing. **d**, cryo tomographic slice of nuclear envelope with inner nuclear membrane evagination depicted as stars (*nup116Δ* 25°C). Labels as in **a**. **e,** Box plot (median, 1st quartile) showing that the number of inner nuclear membrane evaginations are significantly higher in the NEs of *nup116Δ* cells at 25°C (28 in 130 cryo tomograms) in comparison to WT envelopes (1 in 230 cryo tomograms); p<0.00001 (Mann Whitney, two-sided).

**Extended Data Fig. 7:**
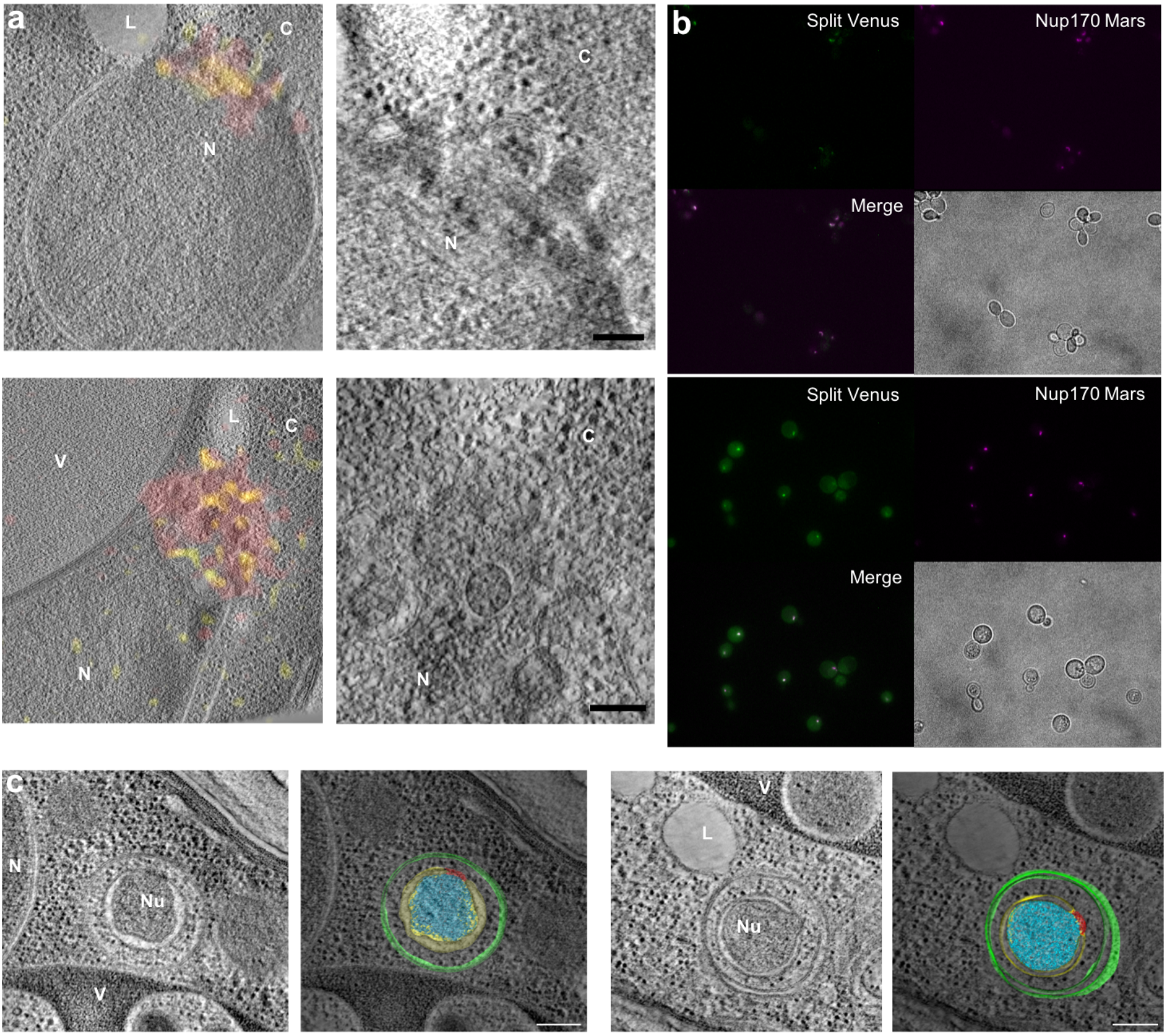
Snapshots of selective autophagy at the nuclear envelope. **a,** Gallery of tomographic slices overlaid with fluorescence images as obtained by on section-CLEM of Nup170-Mars, Nup159-Atg8-SplitVenus cells. The Nup159-Atg8 interaction is observed as the yellow SplitVenus signal, Nup170-Mars is shown in red. In the right panel, zoomed views of the same tomographic slices are shown, emphasizing the clustering of herniae and nuclear vesicles arising through further membrane remodeling (See also Fig. 3 c-d and Supplementary Video 1). N is nucleus, V is vacuole, L is lipid droplet, C is cytoplasm. Scale bar: 100 nm. **b**, Fluorescence and phase contrast microscopy images of Nup170-Mars (magenta), Nup159-Atg8-SplitVenus (green) cells before (top) and after 5,5h of nitrogen starvation (bottom). Upon starvation, Nup159-Atg8-SplitVenus signal increases and co-localizes with Nup170-Mars (white dots). **c,** tomographic slice (left) as well as the corresponding segmentation and isosurface rendering (right) of NPC-containing nuclear vesicles (~340 nm in diameter) observed in the cytoplasm. They are surrounded by cytosol content (ribosomes) and another two membranes (~540 nm in diameter). *Sce atg15D*cells were starved for ~24 h to enrich NPC autophagy events. NPCs in red, Nuclear content (Nu) in cyan, autophagosomal double membranes in green. Scale bar: 200 nm.

**Extended Data Fig. 8:**
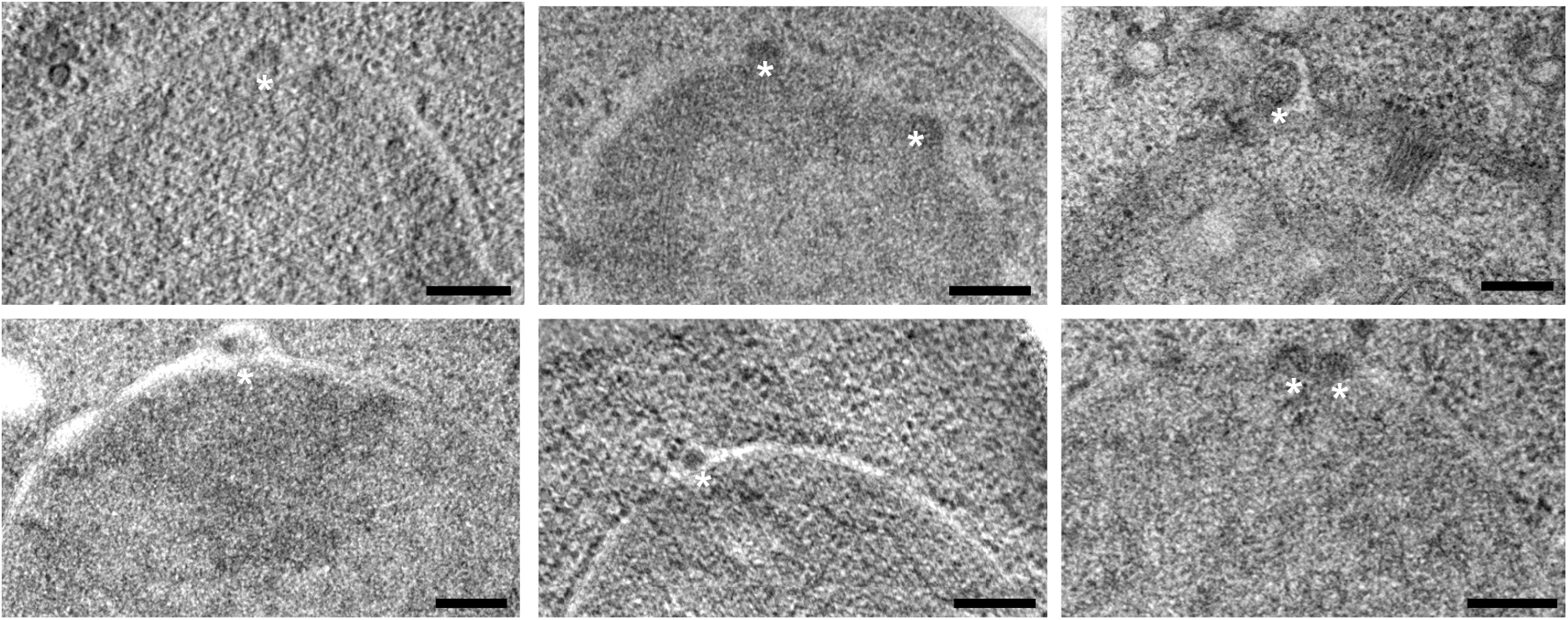
Herniations under *atg8Δ* conditions after 5h of nitrogen starvation. Small herniations appears in the nuclear envelope and are marked with *. Statistics in Figure 3b. Scale bar: 200 nm.

**Extended Data Fig. 9:**
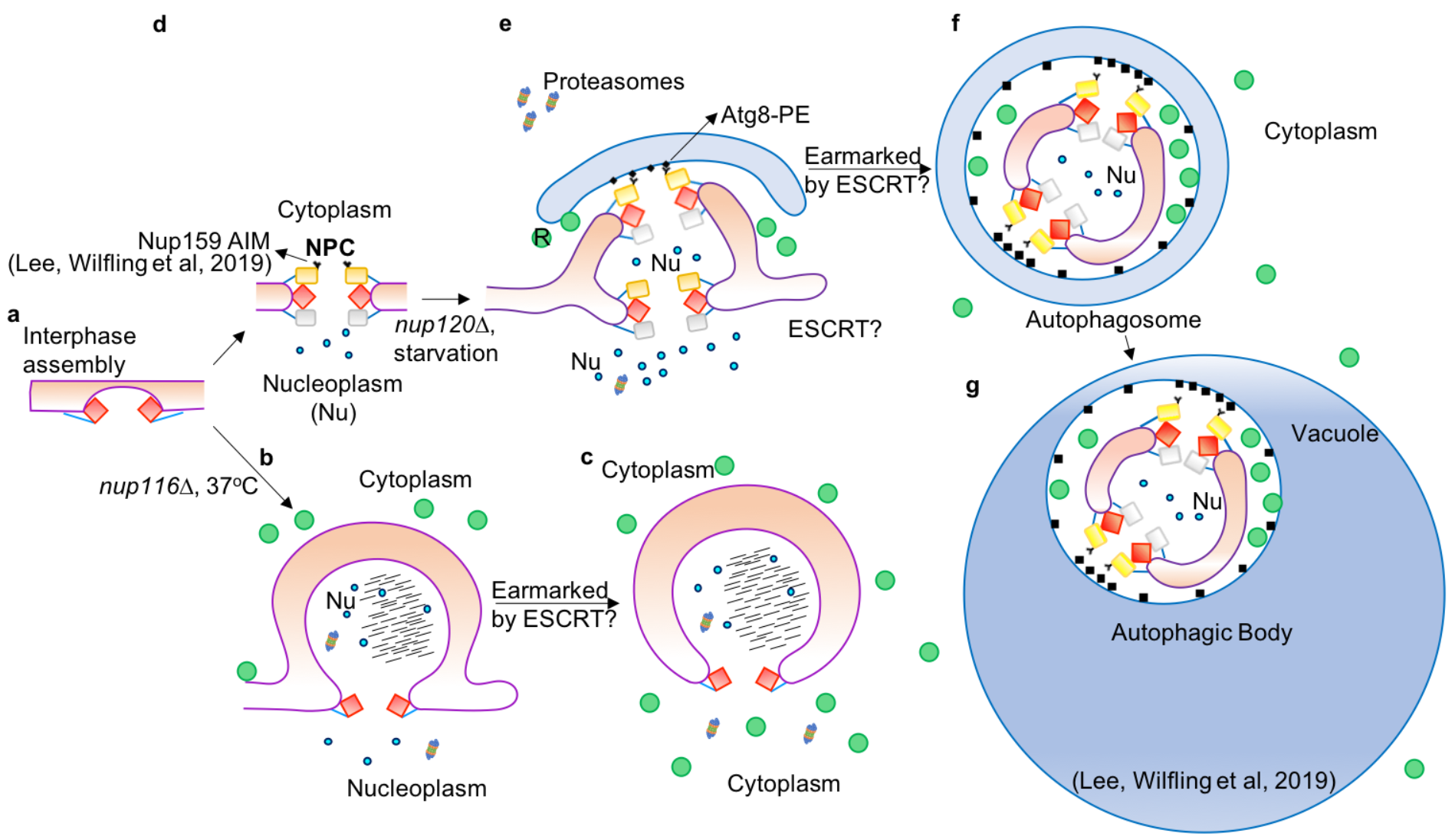
Cartoon summarizing the results from this paper and the accompanying paper Lee et al. Color code as in Fig. 4.

## Legends for Supplementary items

**Video Supplementary 1: 3D cryo-CLEM tomogram of Nup159-Atg8 interaction**. The video shows how clustered NPCs depicted in red are surrounded by double membranes depicted in yellow after 5,5h starvation. Fig. 3b for statistics and 3d for scale bar.

**Video Supplementary 2: Cryo electron tomogram of *nup116Δ* strain #4 after shifting 5h to 37°C shows budded herniae**. NPCs at the herniae are red, membranes are pink. Scale bar: 100 nm.

**Extended Data Table 1: *S. cerevisiae* strains used in this study**.

Source Data 1: Comparison of Cryo-EM maps of the *Sc*NPC. Zip file for reviewers containing the in cell cryo-EM map of the *Sc*NPC (this study), the cryo-EM map (EMD-7321) of detergent extracted NPCs previously obtained by Kim et al. and a Chimera session that displays both drawn to scale at suggested contour level (in cell cryo-EM map in grey, EMD-7321 in cyan).

## Methods

### Yeast strains, growth conditions, fluorescence imaging

Strains used in this study are listed in Extended Data Table 1. Fresh cells were grown to mid-log phase in YPAD (1% Yeast Extract + 2% Peptone (YP), adenine (A), 2% Dextrose (D)). Nitrogen starvation was carried out by switching cells grown at OD_600_ ~0.5 with YPAD into SD-N medium (synthetic minimal medium lacking nitrogen; 0.17% yeast nitrogen base, without aminoacids and ammonium sulfate, supplemented with 2% glucose) for indicated times. Fluorescent images of unfixed yeast cells upon starvation (strains #2, 3) were acquired using a Zeiss Axio Observer. Colocalization figure was made in FIJI^38^.

### Correlative fluorescence and electron tomography in plastic sections

CLEM analysis was conducted as previously described^39,40^. *S cerevisiae (Sc)* cells strain #2 after ~5h of starvation were high pressure frozen (HPM010, AbraFluid) and freeze substituted (EM-AFS2, Leica) with 0.1% uranyl acetate in acetone for 55h at −90°C. The temperature was then raised to −45°C at 3.5°C/h and samples were further incubated for 5h. After rinsing in acetone, the samples were infiltrated in Lowicryl HM20 and the resin was polymerized under UV light. 300 nm sections were cut with a microtome (EM UC7, Leica) and placed on carbon coated 200 mesh copper grids (S160, Plano).

The fluorescence microscopy (FM) imaging of the sections was carried out as previously described^36,40^ using a widefield fluorescence microscope (Olympus IX81) equipped with an Olympus PlanApo 100X 1.40 NA oil immersion objective and a CCD camera (Orca-ER; Hamamatsu Photonics).

After fluorescence imaging, grids were poststained with uranyl acetate and lead citrate. 15 nm protein A-coupled gold beads were also added as fiducial markers used for overlaying low-mag with high-mag tomograms. 60 ° to 60 ° tilt series of the cells of interest were acquired semi-automatically on a Tecnai F30 (Thermofisher, FEI) at 300 kV with Serial-EM^41^ at 20000x and at 4700x to facilitate ease of correlation.

Tomograms were reconstructed automatically with IMOD batchruntomo^42^. Tilts were aligned by patch tracking. 3DMOD software was used for manual segmentation of the tomograms and for making the videos. Overlays of fluorescence spots and tomograms was performed with ec-CLEM Plugin^43^ in ICY^44^ by clicking manually on corresponding pairs of notable features in the two imaging modalities.

### 2D electron microscopy

Strains #1, 2, 3, 5 upon starvation were high pressure frozen as described above and freeze substituted with 0.2% uranyl acetate 0.1% glutaraldehyde in acetone for 60h at −90°C. The temperature was then raised to −45°C at 3°C/h and samples were further incubated for 9h. After rinsing in acetone, the samples were infiltrated in Lowicryl HM20 resin and the resin was polymerized under UV light. Acquisition was performed with a Tecnai F30 (Thermofisher, FEI) at 300 kV. Herniation counting was performed using 2D electron microscopy of 70-100 nm sections and 3D electron microscopy using 300 nm sections. Boxplots were done with BoxPlotR (http://shiny.chemgrid.org/boxplotr/) using around 150 nuclei counted per each strain or condition.

### Cryo-FIB milling and cryo-CLEM

*Sc* wild type strain #1 and knock-out #4 were grown in YPAD liquid medium to an OD_600_ of 0.2 at 30°C, OD 0.5 for correlative studies using strain #2. Per grid, 3.5ul of cell suspension were applied on 200-mesh copper grids coated with R 2/1 holey carbon or SiO2 films (Quantifoil Micro Tools GmbH) and plunge frozen in liquid ethane at ~-186°C using a Leica EM GP grid plunger. The blotting chamber conditions were adjusted to 30°C, 95% humidity and 1,5-2 seconds blotting time. For cryo-CLEM, Crimson FluoSpheres™ Carboxylate-Modified Microspheres, 1.0 μm, crimson fluorescent (625/645), were washed in 1×PBS and added to the cell suspension at 1:20 dilution, and 1 second blotting time was used.

The frozen grids were fixed in modified autogrids to allow milling at a shallow angle^45^ and transferred into an Aquilos (cryo-FIB-SEM dual beam, Thermofisher). For correlative studies, the clipped grids were imaged on a prototype Leica cryo-confocal microscope based on Leica TCS SP8 CFS equipped with a Cryo Stage (similar to the commercially available EM Cryo CLEM Widefield system). Imaging was performed using a 50× objective, NA 0.90, 552 nm laser excitation, and detecting simultaneously at 560–620 nm and 735–740 nm.

In the Aquilos, samples were sputter-coated with inorganic platinum and coated with an organometallic protective platinum layer using the Aquilos gas injection system (GIS)^46^. Lamellae were produced using the Gallium ion beam at 30 kV and stage tilt angles of 17°-19° by milling in two parallel rectangular patterns. The lamella preparation was conducted in a step-wise fashion, gradually reducing the current of the ion beam until the final polishing of a thin slab of biological material of around 150-250 nm.

For correlative studies, beads (5 to 10) were picked in the squares of interested and overlaid with fluorescence signals coming from the confocal stacks using 3DCT in both the electron and ion beam images^37^. After choosing the signal of interest in the confocal stacks, 3DCT provides the position to place the milling patterns in x, y and zed. Milling is then performed as described above. After polishing, the signal of interest is retained in the lamellae provided that the milling did not result in deformation of the carbon film.

Autogrids with lamellae were unloaded and placed in storage boxes. In some cases, a further inorganic Pt layer was added for reducing charging during TEM imaging.

### Cryo-electron tomography data collection and processing

Grids with lamellae were loaded into the Krios cassette. Cryo-electron tomographic tilt series (TS) were acquired on a Titan Krios (Thermofisher FEI) operating at 300 kV equipped with a Gatan K2 summit direct electron detector and energy filter. The autogrids were carefully loaded with the lamella orientation perpendicular to the tilt-axis of the microscope prior to TS acquisition.

Lamellae were mapped at low mag (2 nm/pix, 30 kV slit) and for correlative studies the maps were overlaid with the electron beam images from the Aquilos using serialEM registration points. Spots of interested were chosen accordingly.

All data was collected using the K2 operating in dose-fractionation mode at 4k x 4k resolution with a nominal pixel size of 0.34 nm. TS collection was automated at using a modified version of the dose-symmetric scheme^47^ taking the lamella pre-tilt into account. Defocus TS were acquired over a tilt range of +65 to −45 for a positive pre-tilt with a tilt increment of 2-3°, a total dose of ~140 e-/A^2^ and a targeted defocus of around 2 to 4.5 μm.

All images were pre-processed and dose-filtered as described in^11^. Tilt-series alignment was performed using IMOD software package 4.9.2.^42^, by patch tracking function on bin4 image stacks. Initial tilt series alignment was manually inspected and improved by removing contours showing large deviations (i.e. a large mean residual error) from the alignment model function. The software package gCTF^48^ was used for CTF estimation. 3dCTF correction and tomogram reconstruction was performed with NovaCTF^49^.

### Subtomogram alignment and averaging

Subtomogram alignment and averaging was performed with slight modifications from a previously described workflow^50^ using the Matlab TOM package re-implemented in C++. In brief: Particle coordinates and initial orientations were manually picked and assigned. Initial NPC alignment was performed on 8 and 4 times binned subtomograms followed by a manual inspection and curation of the initial alignment for each particle. From the whole aligned NPCs all 8 spokes were assigned according to an 8-fold rotational symmetry and the subunits were extracted from the tomograms removing subunits which are located outside of the lamellae. Subunits were further aligned. The alignment was once more manually inspected, and all misaligned subunits were removed. Final alignments were done binning two times the subtomograms using focused masks on the cytoplasmic, inner and nucleoplasmic ring of the NPC.

In the *S. cerevisiae* wild type data set ~500 NPC (4000 asymmetric units) were initially picked from ~230 tomograms, ~200 NPCs (1600 asymmetric units) from 120 tomos for the *nup116Δ* strain at permissive temperature and ~40 NPCs from 40 tomos for the *nup116Δ* strain at not-permissive temperature (37°C). The particles were split in two half datasets for gold-standard processing. After alignment and manual curation 2000 subunits were used in the final average for the WT NPC and ~800 for the *nup116Δ* at permissive temperature and 320 for the *nup116Δ* strain at 37°C. For the latter NPCs filtered volumes were used for alignment and weighted back projection volumes to get the final structure. Gold standard Fourier shell correlation (FSC) was calculated with Fourier Shell Correlation Server (www.ebi.ac.uk/pdbe/emdb/validation/fsc/results/) with half-maps as inputs. Local resolution was calculated with Resmap^51^, B-factor sharpening was estimated empirically as in^9^ being between 2000 Å^2^ for the inner ring and 3000 Å^2^ for the cytoplasmic ring. For counting the number of inner nuclear membrane evaginations (Extended Data Fig. 6d-e), all the WT tomograms and ~120 tomograms from *nup116Δ* cells at permissive temperature were used.

### Systematic fitting of *Sc*NPC components to the cryo-ET map

An unbiased systematic (global) fitting approach was performed using structural models of various *Sc*NPC subcomplexes derived from previously published structures^3,13^. All structural models were low-pass filtered to 40 Å prior the fitting. The resulting model maps were then independently fitted into the *Sc*NPC cryo-EM map using global fitting as implemented in UCSF Chimera^52^. The fitting of the Y complex structures and Nup159 complex was performed for the isolated maps of the cytoplasmic and nuclear rings. The inner ring model was fitted to the IR map. The nuclear envelope density was erased prior to the fitting. The regions of the nuclear envelope distant from apparent contact points between membrane and protein densities were erased prior to the fitting to eliminate fits significantly overlapping with the membrane. All fitting runs were performed using 100,000 random initial placements and the requirement of at least 30% of the model map to be covered by the *Sc*NPC density envelope defined at low threshold. For each fitted model, this procedure yielded between 500-17,000 fits after clustering.

For fitting the filtered atomic models, the cross-correlation about the mean (cam score, equivalent to Pearson correlation) score from UCSF Chimera was used as a fitting metric as in our previous work^9,10,11,12^.

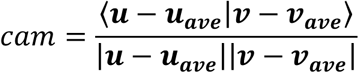

where **u_*ave*_** is a vector with all components equal to the average of the components of **u** and **v_*ave*_** is defined analogously.

The negative stain map of the Nup159 complex was fitted using Chimera’s cross-correlation about zero (equivalent to cosine similarity).

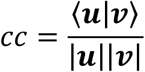

This score was used instead of the cam score because the cam score tests the linear dependence of the two fitted maps, which is not the case when fitting negative stain maps, which contain the signal of the electron density at the surface of the complex.

To confirm this fit, the Nup159 complex was also fitted using the colors program from the Situs package^54^ with settings appropriate for low resolution negative stain maps (i.e. using Laplacian filtering that emphasizes on contour matching over interior volume matching).

For each fitting run, the statistical significance of the fits was assessed as a p-value calculated from the normalized cross-correlation scores. To calculate the p-values, the cross-correlation scores were first transformed to z-scores (Fisher’s z-transform) and centered, from which two-sided p-values were computed using standard deviation derived from an empirical null distribution (derived from all obtained fits and fitted using fdrtool^55^ R-package). All p-values were corrected for multiple testing using Benjamini-Hochberg procedure. Figures were made using UCSF Chimera^52^ and Xlink Analyzer^56^.

### Integrative modeling

To build the integrative model of *Sc*NPC, the IR model from^13^ was fitted as a rigid body using the procedure described above. The models of Y-complex fitted into CR and NR could not be obtained by rigid body fitting of the published models^13^. Therefore, we have divided the available crystal structures and homology models^13^ into six smaller rigid bodies (with cut points corresponding to boundaries of published crystal structures of Y-subcomplexes) and fitted them simultaneously into the cryo-EM maps of the CR and NR using the integrative modeling procedure implemented with using Integrative Modeling Platform^57^ version 2.9.0 as described previously by^11,53^. Firstly, each of the rigid bodies has been independently fitted to the EM map using UCSF Chimera^52^ as described above (structural models were low-pass filtered to 40 Å) to generate libraries of alternative fits for each rigid body. Then, we generated configurations of all the rigid bodies by recombining the above fits using simulated annealing Monte Carlo optimization. Each configuration was generated by an independent Monte Carlo optimization comprising 30,000 steps resulting in total 20,000 models and scored. The scoring function for the optimization was a linear combination of the normalized EM cross-correlation scores of the precalculated domain fits, domain connectivity restraint, a term preventing overlap of the Y components with the nuclear envelope and the assigned Nup159 complex density) and clash score (see ref.^11^ for the implementation details). The structures were simultaneously represented at two resolutions: in Cα-only representation and a coarse-grained representation, in which each 10-residue stretch was converted to a bead. The 10-residue bead representation was used for the clash score to increase computational efficiency, the Cα-only representation was used for crosslinking and domain connectivity restraints. Since the EM restraint was derived from the original EM fits generated with UCSF Chimera, it was derived from the full atom representation. The final models for visualization were selected as the top scoring model.

## Acknowledgments

We thank members of the Beck Lab, Mahamid Lab and EMBL’s electron microscopy core facility for invaluable input and support. We thank Leica for collaboration to develop the prototype cryo-confocal. We thank Amparo-Andres Pons, David Thaller, Patrick Lusk and Ed Hurt for critical discussions and for providing relevant strains. We thank Jiameng Sun for help in tomographic segmentation. The computational part of this work has been done on the High-Performance Cluster of the EMBL, supported by the EMBL Heidelberg IT Services. This work was supported by ERC Complex Assembly (MB and MA); MA was funded by an EMBO a long-term fellowship (ALTF-1389–2016); JM received funding from the European Research Council (ERC 3DCellPhase^-^ 760067); KHF is supported by a fellowship from the EMBL Interdisciplinary (EI3POD) programme under Marie Skłodowska-Curie Actions COFUND (664726). MB acknowledges funding by EMBL, the Max Planck Society and the European Research Council (ComplexAssembly 724349).

## Author Contributions

MA, CEZ conceived the project, designed experiments, collected data, analyzed data, wrote the manuscript. VR analyzed data, performed computational analysis, wrote the manuscript. FW, PR designed experiments, collected data, analyzed data. HKHF, CWL, XZ collected data, analyzed data. WH collected data. BT analyzed data. KK performed computational analysis. CWM, JM, YS supervised the project. BP, JK, MB conceived the project, designed experiments, supervised the project, wrote the manuscript.

## Depositions

The EM maps associated with this manuscript will be deposited in Electron Microscopy Data Bank upon publication. The integrative model of *Sc*NPC will be deposited in PDB-dev (https://pdb-dev.wwpdb.org/). An in cell cryo-EM map of the *Sc*NPC is attached for the review process (Source Data 1).

